# Small RNA *ASpks2* Promote *Mycobacterium tuberculosis* Survival in Macrophages *via* Targeting Polyketide Synthase 2

**DOI:** 10.1101/2025.06.14.659662

**Authors:** Qian Wu, Mingkun Yang, Lijun Bi, Ying Chen, Wei Yu, Feng Ge

## Abstract

*Mycobacterium tuberculosis* is the causative pathogen of human tuberculosis and a leading cause of death worldwide attributed to a single infectious agent. While small non-coding RNAs (sRNAs) have emerged as key regulators of bacterial pathogenicity, their specific roles and mechanisms in *M. tuberculosis* remain poorly understood. Here, we employed label-free quantitative proteomics, parallel reaction monitoring, and sRNA-seq analyses to identify proteomic differences between the virulent H37Rv and attenuated H37Ra strains. Bioinformatic analysis revealed significant enrichment of differentially expressed proteins involved in lipid metabolism and fatty acid biosynthesis, key pathways linked to *M. tuberculosis* virulence. We identified a novel sRNA, *ASpks2*, which was significantly downregulated in H37Rv. Functional validation demonstrated that *ASpks2* directly targets the polyketide synthase 2 (*pks2*), modulating its expression to enhance *M. tuberculosis* survival in human macrophages THP-1 cells. By correlating the omics data with functional studies, this study identified a novel sRNA and its regulatory network in *M. tuberculosis*, which provides novel insight into the molecular pathogenesis of *M. tuberculosis* and may serve as a basis for the development of targeted therapies.

## Introduction

Tuberculosis (TB) persists as a formidable adversary to global health, with 10.6 million new cases reported in 2022 and an incidence rate of 133 per 100,000 individuals. TB claimed 1.3 million lives, making it the second leading cause of death from a single infectious pathogen, subordinate only to COVID-19, and approaching a twofold increment of HIV/AIDS ^1^. *Mycobacterium tuberculosis* (*M. tuberculosis*) primarily infected the lungs and current TB control strategies face significant limitations. The Bacillus Calmette-Guérin (BCG) vaccine, while widely used, offers limited and inconsistent protection, particularly in adults ^2^. Standard treatment requires a six-month or longer regimen with multiple antibiotics, posing risks of hepatotoxicity and drug resistance. Alarmingly, only three new drugs targeting drug-resistant tuberculosis have been developed in the past 50 years, underscoring the urgent need for innovative research into the pathogenic mechanisms of *M. tuberculosis* ^3^.

The virulent *M. tuberculosis* H37Rv and its attenuated H37Ra strain serve as pivotal models for studying the molecular basis of *M. tuberculosis* pathogenicity. Genomic comparisons have identified variations in sequences such as IS6110, the RD1 region, and the PE/PPE gene families as the primary drivers of genetic divergence ^4–7^. Jia et al. identified specific single nucleotide polymorphisms (SNPs) associated with attenuated virulence, including S219L in PhoP, A219E in MazG, and the newly identified I228M in EspK ^8^. Transcriptome analyses revealed that 22 genes were consistently upregulated in H37Rv compared to H37Ra, with significant enrichment in lipid metabolism and cell membrane function pathways ^9^. Notably, the expression of RD1 genes cfp10 and esat6 is notably absent in the less virulent strain ^7^. Proteomic studies have identified differential protein abundance between H37Rv and H37Ra, particularly in fatty acid biosynthesis and bacterial secretion systems ^10, 11^. Hiwa Mamatlen et al. have further characterized a set of 19 membrane and lipoproteins with higher abundance in H37Rv, and 10 proteins with higher abundance in H37Ra ^12^. Despite these findings, the precise mechanisms driving these differences remain unclear.

Small RNAs (sRNAs), non-coding RNA ranging from 50 to 500 nucleotides, have emerged as pivotal regulators in bacterial pathogenicity. These molecules modulate mRNA translation and stability by base-pairing with target mRNAs, enabling bacteria to adapt to stress and host environments ^13, 14^. Advances in RNA sequencing have identified numerous sRNAs in *M. tuberculosis* ^15–18^, many of which display distinct expression patterns under stress conditions, such as exposure to patient serum or plasma, macrophage infection, oxidative stress, pH fluctuations, hypoxia, iron limitation, and starvation. These findings suggest that sRNAs are implicated in processes like latency, growth, infection, and regulatory adjustments ^19–25^, and they may play pivotal roles in regulating critical biological processes in *M. tuberculosis*. For instance, *Mcr7* modulates the twin-arginine translocation (Tat) protein secretion system by targeting *tatC* mRNA ^26^, while *MrsI* (*ncRv11846*) influences iron homeostasis by interacting *bfrA* ^27^. Additionally, 47 genes have been delineated as potential direct targets of the 6C sRNA, with 15 of these validated using *in vivo* translational lacZ fusion systems ^28^. Despite these advances, the differential expression of sRNAs and their regulatory networks in the virulent H37Rv and attenuated H37Ra strains remains unexplored.

In this study, we integrate label-free quantitative proteomics, parallel reaction monitoring (PRM), and sRNA-seq analyses to unravel the molecular differences between H37Rv and H37Ra. A novel sRNA, *ASpks2*, was identified as a key regulator of the polyketide synthase 2 (*pks2*) pathway, a critical determinant of lipid metabolism and virulence. By establishing the regulatory relationship between *ASpks2* and *pks2*, this research provides new insights into the molecular mechanisms driving *M. tuberculosis* survival in macrophages and offers potential avenues for therapeutic intervention.

## EXPERIMENTAL PROCEDURES

### Biosafety Notes

*M. tuberculosis* H37Rv was handled in a biosafety level 3 (BSL-3) laboratory under stringent biosafety protocols to minimize exposure risks. The attenuated H37Ra strain was handled in a BSL-2 environment ^29^. World Health Organization (WHO) Biosafety Manual for Tuberculosis Laboratories provides a series of safety measures to ensure that the risk of accidental transmission is minimized when conducting experiments related to *M. tuberculosis* (including H37Rv and H37Ra) ^30^.

### Bacterial strains and culture conditions

*M. tuberculosis* H37Rv and H37Ra were cultured in Middlebrook 7H9 medium supplemented with 10% (v/v) OADC (oleic acid, albumin, dextrose, catalase) enrichment, 0.5% glycerol, and 0.05% Tween 80. For plasmid-containing strains, 20 μg/mL kanamycin was added. Cultures were grown at 37°C under static conditions until the logarithmic phase (OD_600_ = 0.7-0.8), at which point cells were harvested for downstream RNA and protein extraction.

### Protein extraction and digestion

Bacterial cells were washed twice with ice-cold PBS buffer, resuspended in ice-cold PBS containing 1 × protease inhibitor mixture (Roche Diagnostics Ltd) and 1 mM PMSF. The mixture was processed in a nucleic acid extractor Fastprep-24 (MP Biomedicals) for 35 s at a speed of 6.5 m/s for 5 cycles as previously described ^31^. Cell lysates were centrifuged at 12000 g for 15 min at 4°C to remove cellular debris. Supernatant was then filtered through the 0.2 μm syringe filters (Merck Millipore) twice to sterilize lysate. The protein concentration was determined by BCA assay (Beyotime) following the manufacturer’s instructions.

Proteins were reduced, alkylated, and trypsin digested as described previously ^32^. Briefly, 100 ug of each group of samples was reduced with 25 mM dithiothreitol (DTT) for 45 min at 37°C and alkylated with 50 mM iodoacetamide (IAM) for 10 min in the dark. Then, the samples were digested with a trypsin to protein mass ratio of 1:100 at 37°C for 24 h, following by the addition of 0.1% (v/v) trifluoroacetic acid (TFA) to terminate the digestion. The digests were finally desalted using self-packed C_18_ STAGE column and dried using a vacuum centrifuge.

### LC-MS/MS analysis

After dissolving with 1% formic acid (FA), peptides were separated on an online nanoEASY1200 HPLC system (Thermo Fisher Scientific) coupled with an Orbitrap Q Exactive HF-X mass spectrometer (Thermo Fisher Scientific) on a 75 μm × 15 cm analytical column (Thermo Fisher Scientific). The nanopump provided a flow-rate of 300 nL/min and was operated under gradient elution conditions, using 0.1% FA as buffer A, and 0.1% FA in 90% ACN as buffer B. Gradient elution was performed according the following scheme: 1 min of gradient from 4%-8% buffer B, 34 min of gradient from 8%-13% buffer B, 26 min of gradient from 13%-18% buffer B, 25 min of gradient from 18%-25% buffer B, 19 min of gradient from 25%-35% buffer B, 10 min of gradient from 35%-40% buffer B, 1 min of gradient from 40%-100% buffer B, and a final 9 min of 100% buffer B to wash the column. Subsequently, the eluted peptides were ionized and detected using the Orbitrap Q Exactive HF-X mass spectrometer (Thermo Fisher Scientific). MS data were collected in data-independent acquisition (DIA) mode using 100 variable windows covering a mass range of 350-1200 at a resolution of 60,000. The normalised AGC target was set to 3E6 and the maximum injection time was 50 ms. Following every survey scan, the m/z range of 400-1000 was acquired at 30,000 resolution with 6 m/z of isolation window for DIAscan. Precursor ions were selected for fragmentation by high-energy collision dissociation with a normalized collision energy of 30%. The normalised AGC target was set to 2E5 and the maximum injection time was set to auto.

### Database search and protein quantification

LC-MS raw files were processed using DIA-NN software (version 1.8.1) ^33^ using a library free workflow against the H37Rv (GCA_000195955.2) and H37Ra (GCA_000016145.1) databases concatenated with common contaminants. “FASTA digest for library free search/library generation” and “Deep learning spectra, RTs and IMs prediction” options were used for precursor ion generation. Two missed cleavages were allowed for trypsin. Carbamidomethylation (Cys) was set as a fixed modification, whereas dynamic modifications were set as oxidation (Met) and acetylation (Protein N-terminal). Maximum number of variable modifications was set to 1. Peptide length was set to 7-30 amino acids, and the precursor charge range was restricted to 1-4. The FDR at the precursor, peptide and protein levels was set to 1%. Other parameters were kept at their default values in DIA-NN software. Quantification was performed by LFQ algorithm. LFQ intensities were extracted from the DIA-NN report file “report.pg_matrix.tsv”. Data analysis and statistical evaluation were done with Microsoft Excel. Only proteins identified by at least two unique peptides were reported. LFQ intensities had to be detected in two biological replicates in one of the groups were included in the relative quantitative analysis. Missing values in the third biological replicate were estimated using the default parameters for normal distribution in Perseus ^34^. The total LFQ intensities of all identified proteins were normalized and the average LFQ intensity for each biological sample group was derived by calculating the arithmetic mean. Statistical assessments were performed using a two-sample Student’s t-test, and differentially expressed proteins (DEPs) were defined as fold change (FC) > 1.5 or < 0.67 with *p*-value < 0.05.

### DIA and parallel reaction monitoring (PRM) experiments

For DIA-based experiments, the digested samples were analyzed on an Ultimate 3000 HPLC system (Dionex) coupled with an Orbitrap Exploris™ 240 mass spectrometer (Thermo Fisher Scientific) on a 75 μm × 15 cm analytical column (Thermo Fisher Scientific). Peptides were eluted using a 135 min gradient of buffer B as follows: 0 to 5 min, 2% solvent B at a flow rate of 400 nL/min; 5 to 5.5 min, 2% solvent B at a flow rate of 300 nL/min; 5.5 to 11 min, 2%-3% solvent B at a flow rate of 300 nL/min; 11 to 40 min, 3%-7.5% solvent B at a flow rate of 300 nL/min; 40 to 102 min, 7.5%-18% solvent B at a flow rate of 300 nL/min; 102 to 117 min, 18%-25% solvent B at a flow rate of 300 nL/min; 117 to 122 min, 25%-33% solvent B at a flow rate of 300 nL/min; 122 to 127 min, 33%-55% solvent B at a flow rate of 300 nL/min; 127 to 128 min, 55%-99% solvent B at a flow rate of 400 nL/min; 128 to 135 min, 99% solvent B at a flow rate of 400 nL/min. The eluted peptides were ionized and detected using the Orbitrap Exploris™ 240 mass spectrometer (Thermo Fisher Scientific). MS data were collected in data-independent acquisition (DIA) mode using 100 variable windows covering a mass range of 350-1200 at a resolution of 120,000. The normalised AGC target was set to 300% and the maximum injection time was 50 ms. Following every survey scan, the m/z range of 400-1000 was acquired at 30,000 resolution with 6 m/z of isolation window for DIAscan. Precursor ions were selected for fragmentation by high-energy collision dissociation with a normalized collision energy of 30%. The normalised AGC target was set to 200% and the maximum injection time was set to 100 ms. All raw files were processed using DIA-NN software (version 1.8.1) with the same parameters as described above (“Database search and protein quantification”). The peptide identification results were imported into Skyline software ^35^ as a spectral library. Based on the spectral library results, the transition lists of the target tryptic peptides were generated and used for PRM analysis.

For PRM experiments, the digested samples were analyzed on an Ultimate 3000 HPLC system (Dionex) coupled with an Orbitrap Exploris™ 240 mass spectrometer (Thermo Fisher Scientific) on a 75 μm × 15 cm analytical column (Thermo Fisher Scientific). MS data collection was performed in PRM acquisition mode with the same gradient elution conditions and MS parameters as described in the DIA-based experiments. All PRM data were processed using Skyline software as previously described ^36^. Briefly, to avoid false peptide identification, 3 transitions per peptide were at least recommended to ensure enough transitions and to maintain the selectivity for reliable quantification. Peaks were manually checked for correct integration, and the area under the curve (AUC) of targeted peptide was obtained from the summation of the AUC for each transition peak. The abundance of each peptide was normalized based on the average abundance of each protein. For protein quantification, the average of the target peptide abundances was used to calculate the fold change between samples.

### Bioinformatics analysis

Gene Ontology (GO) and Kyoto Encyclopedia of Genes and Genomes (KEGG) pathway enrichment analyses were performed using STRING and DAVID databases ^37–39^. The GO terms and KEGG pathways were considered significantly enriched when they met a *p*-value < 0.05. Using the TubercuList database for functional category ^40^. Genome analysis was performed using TBtools software ^41^. The heatmaps were generated using R software (4.2.1) and the ComplexHeatmap package (2.13.1). Protein-protein interaction (PPI) network analysis was performed using the STRING database. The network was visualized by Cytoscape software (v3.9.1) and further analyzed for densely connected regions using the cytoHubba plugin in Cytoscape ^42^. Correlation values were calculated using the Pearson correlation coefficient test.

### RNA extraction and sRNA-seq

The small RNA (sRNA) sequencing data for the H37Rv strain were retrieved from previous study ^16^. For H37Ra, we collected the bacterial cells that grown in the same cultivation condition with H37Rv strain. The cells were lysed using a bead mill, and the resulting supernatant was subjected to centrifugation. Further purification involved two chloroform extractions, isopropanol precipitation, and washing with 80% ethanol to ensure high RNA purity. The extracted RNA was treated with 10 U DNase I (Fermentas) at 37°C for 30 min to prevent genomic DNA contamination, followed by additional purification steps. A total of 10 μg of RNA was further processed to deplete rRNA using the MICROBExpress Kit (Ambion) according to the manufacturer’s instructions. The RNA was fractionated via urea polyacrylamide gel electrophoresis into three size classes: Fraction 1 (18-40 nt), Fraction 2 (40-80 nt), and Fraction 3 (80-140 nt). The remaining sample, consisting of rRNA-depleted RNA, was fragmented to ∼140 nt using divalent cations at high temperatures. Then, strand-specific cDNA libraries were prepared following the Illumina TruSeq protocol, as previously described ^43^. For fractions 1 and 2 samples, cDNA constructs were prepared by reverse transcription followed by low-cycle PCR amplification. PCR products were collected by gel purification and sequenced on the Illumina Genome Analyzer II (Illumina, San Diego, USA) platform. For fractions 3 and 4 samples, RNA libraries were constructed using the TruSeq mRNA kit (Illumina) according to the manufacturer’s instructions and sequencing was performed on an Illumina Genome Analyzer II (Illumina, San Diego, USA) platform.

Raw reads were processed as described by Wang et al ^16^. Briefly, low-quality reads and adapter sequences were removed using Trimmomatic tools (version 0.32) ^44^ with the following criteria: 1) Removal of adapter and primer-matched sequences; 2) Trimming leading and trailing bases with a Phred33 score <3; 3) Applying a sliding window of 4 bases, cutting reads when the average base quality dropped below 15; 4) Discarding reads shorter than 16 bases after trimming. The clean reads were then aligned to the *M. tuberculosis* H37Rv reference genome (NC_000962.3) or H37Ra reference genome (NC_009525.1) using Bowtie2 ^45^ with default parameters. The strand-specific read coverage at each genomic base were generated using SAMtools^46^ and BEDTools ^47^. Data were visualized in the Integrated Genome Browser (IGB) ^48^ to assess sequence coverage at each genomic position. In addition, BLAST was used to compare sequences and match sRNAs between H37Rv and H37Ra strains ^49^.

### Production of Polyclonal Antibodies

Anti-Hsp (Heat shock protein Hsp), Fgd1 (F420-dependent glucose-6-phosphate dehydrogenase Fgd1), MtrA (Two component sensory transduction transcriptional regulatory protein MtrA), Pks2 (Polyketide synthase Pks2), FadD31 (Probable acyl-CoA ligase FadD31), Pks13 (Polyketide synthase Pks13), and AtpF (Probable ATP synthase B chain AtpF) polyclonal antibodies were produced and purified via affinity chromatography by ABclonal (Wuhan, China). Briefly, the full-length or C-terminal region cDNA of H or AtpF was amplified, PCR products were cloned into the pGEX-4T-1 expression vector (Amersham Pharmacia Biotech), and the resulting plasmid was transformed into E. coli strain BL21 (DE3) for overexpression of these proteins. Cells growing logarithmically were treated with 1 mM isopropyl-β-D-thiogalactopyranoside (IPTG) for 4 h at 30°C. The fusion proteins were then purified by GST-tag affinity chromatography. Following purification of these antigens, immunization and sampling of the anti-sera from rabbit were performed by ABclonal (Wuhan, China), according to standard operating procedures. The specificity of the generated antibodies was determined by the manufacturer using ELISA and Western blotting.

### Western blotting

Equal amounts of proteins from H37Rv and H37Ra strains were separated on 10% or 15% SDS-PAGE gels. Proteins were stained with Coomassie Brilliant Blue R250 or transferred to polyvinylidene fluoride (PVDF) membranes (GE Healthcare). Membranes were blocked with 5% nonfat milk for 2 h, and incubated over-night with corresponding protein-specific antibodies (1:1000 dilution). After washing, membranes were incubated with a 1:3000 dilution of peroxidase-conjugated anti-rabbit IgG (Promega) for 1 h at room temperature. Chemiluminescence was detected by using the SuperSignal® West Pico Chemiluminescent Substrate (Thermo Fisher Scientific). Immunoblots were performed in three independent experiments and bands of interest analyzed by ImageJ were expressed as mean ± SD.

### Northern blot hybridization

Total RNA (10 μg) from *M. tuberculosis* H37Rv or H37Ra and a low range ssRNA Ladder (New England Biolabs) were denatured and separated on 8% polyacrylamide gels containing 8 M urea. The separated RNAs were electroblotted onto a Hybond-N+ membrane (Amersham) and cross-linked using short ultraviolet (UV) light. Specific DNA oligonucleotide probes for each candidate sRNAs were radioactively labeled [γ-32P] ATP. Hybridization signals were exposed to a phosphor screen or determined by autoradiography on hyperfilms (Amersham).

### Rapid amplification of cDNA ends (RACE)

For 3’ RACE, a DNA adapter was ligated to total RNA from *M. tuberculosis* H37Rv or H37Ra, Reverse transcription was performed using 50 pmol of a single primer complementary to the DNA adapter. A no-reverse transcriptase control without Superscript III (Invitrogen) was included. PCR was performed using a forward primer specific to target sRNAs and reverse primer specific to the 3’ RACE adapter. PCR products were separated on 8% DNA polyacrylamide gel and bands of interest were excised, cloned and sequenced.

### Construction of overexpression and knockdown strains

The vector was prepared as previously described by replacing the Hsp60 promoter in the pMV261 vector with the RrnB promoter ^15^. Using the primers listed in **Supplementary Table 1**, the insert fragments of sRNA were generated by PCR and then cloned into the modified vector. The resultant transformants were selected on medium containing 20 μg/mL kanamycin. Finally, the correctly sequenced plasmid was transformed into *M. tuberculosis* H37Rv by electroporation.

### THP-1 monocyte cell infection

THP-1 cells were grown in RPMI 1640 supplemented with 10% heat-inactivated fetal bovine serum (Gibco). THP-1 cells were inoculated in 24-well plates at 1×10^6^ cells per well and differentiated in the presence of 0.1 μg/ml PMA (Phorbol-12-myristate-13-acetate) (Sigma) for 2 day. Following PBS washes, cells were infected with *M. tuberculosis* strains at a multiplicity of infection (MOI) of 10. After incubation at 37°C for 4 h, extracellular bacteria were removed by washing with PBS, and THP-1 cells were incubated further. At designated time points, macrophages were lysed in 0.1% SDS, and lysates were plated on 7H10 agar for colony-forming unit (CFU) counts. Macrophage viability was assessed using the trypan blue dye exclusion method. For RNA extraction, 50 ml GTC lysis buffer ^50^ was added to ensure complete macrophage lysis. Bacterial pellet was collected by centrifugation, and RNA from the bacteria was extracted as described above.

### Statistical analysis

All experiments were performed in triplicate, and data were analyzed using two-tailed Student’s *t*-tests, with *p* < 0.05 considered significant.

### Experimental design and statistical rationale

To investigate the differences between the virulent *M. tuberculosis* H37Rv strain and the attenuated H37Ra strain, we compared their genomes, morphological features, biochemical responses, and infection kinetics. These preliminary analyses highlighted differences between the two strains. To further explore their divergence in virulence, a label-free quantitative proteomics analysis was performed by comparing H37Rv and H37Ra strains in three biological replicates. All raw files were searched against the H37Rv (GCA_000195955.2) and H37Ra (GCA_000016145.1) databases using the DIA-NN software (version 1.8.1), and protein quantification was based on LFQ intensity. The FDR thresholds for proteins and peptides were set to a maximum of 1%. Data analysis and statistical evaluation were completed using Microsoft Excel. Statistical assessments were performed using a two-sample Student’s t-test, and differentially expressed proteins (DEPs) were defined as FC > 1.5 or < 0.67 with *p*-value < 0.05. To validate the proteomic data, we performed Parallel Reaction Monitoring (PRM) analysis and western blotting. All PRM data were processed using Skyline software. For peptide identification, three transitions per peptide were at least recommended to ensure enough transitions and to maintain the selectivity for reliable quantification. The average of the target peptide abundances was finally used to calculate the fold change between samples. For sRNA identification, we performed sRNA-seq on both H37Rv and H37Ra strains, and the identified sRNA sequences were compared using the native BLAST tool. To validate the functional role of the novel sRNA, *ASpks2*, we constructed OE-*ASpks2* (overexpression) and KD-*ASpks2* (knockdown) strains, which were confirmed via qRT-PCR and northern blotting. These strains, along with a negative control (NC), were assessed for in vitro growth capacity, in vivo infection capacity, and bacterial survival within host cells. For protein functional annotation, Gene Ontology (GO) and Kyoto Encyclopedia of Genes and Genomes (KEGG) annotation of identified proteins was performed using eggNOG-mapper and TubercuList database. Genome analysis was performed using TBtools software. The heatmaps were generated using R software (4.2.1) and the ComplexHeatmap package (2.13.1). Protein-protein interaction (PPI) network analysis was performed using the STRING database, and the network was visualized by Cytoscape software (v3.9.1). Correlation values were calculated using the Pearson correlation coefficient test. Experimental data are presented as mean ± standard deviation (SD), where *p* < 0.05 were recognized as statistically significant. All experiments were repeated three times to ensure stability of the results and the data from both groups were analyzed comparatively using two-sided Student’s t-test.

## RESULTS

### Genotypic and phenotypic characterization of *M. tuberculosis* H37Rv and H37Ra

*M. tuberculosis* H37Rv, the virulent reference strain, and H37Ra, its attenuated counterpart, serve as pivotal models for dissecting *M. tuberculosis* virulence mechanisms. The attenuation of H37Ra results from specific genomic mutations accumulated during in vitro passage, providing a unique opportunity to study the molecular basis of pathogenicity. Comparative genomic analysis revealed a high degree of similarity between H37Rv and H37Ra genomes, with key distinctions including 53 insertions, 21 deletions, and 76 strain-specific single nucleotide variants (SNVs) in H37Ra (**Fig. 1A**) ^5^. It was reported that the H37Ra genome has an insertion of approximately 8000 base pairs between the 5’ end of the *plcD* gene and IS6110 (RvD2 region), which is strong evidence for distinguishing between H37Rv and H37Ra strains. PCR amplification using primers designed for RvD2 confirmed its presence in H37Ra but not in H37Rv (**Fig. 1B**). Morphologically, H37Rv formed serpentine, cord-like structures with broad surface spreading outward in a thin layer, whereas H37Ra exhibited vertical growth with limited surface expansion (**Fig. 1C**). Under alkaline conditions, H37Rv is capable of staining neutral red dye red, unlike H37Ra. We conducted a comparative analysis of their characteristics *in vitro* culture and discovered that H37Rv exhibited a faster replication rate in 7H9 liquid culture than H37Ra (**Fig. S1**). Additionally, we monitored their proliferation within macrophage THP-1 cells and ascertained that the H37Rv strain could trigger the necrosis of macrophages, significantly reducing the count of viable cells (**Fig. 1D and S2**). During this process, a considerable amount of H37Rv was released from the cell interior, resulting in a significant reduction in the amount of H37Rv inside the cell (**Fig. 1E**).

**Figure 1.**
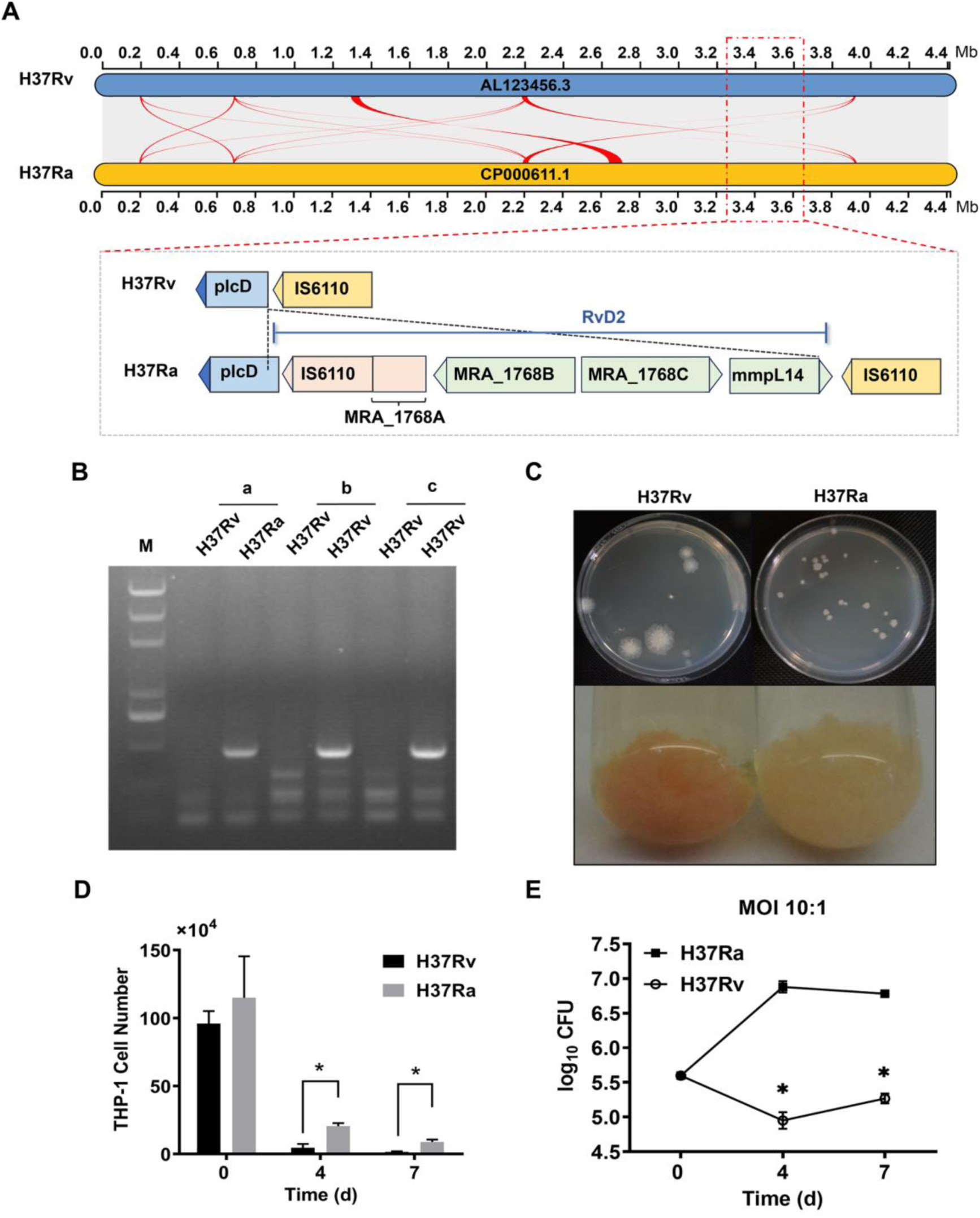
Phenotypic identification of *M. tuberculosis* H37Rv and H37Ra. *A*, comparison of Progressive Mauve analysis between H37Rv and H37Ra genomes. *B*, PCR analysis RvD2 region in H37Rv and H37Ra. Three fragments about 400 bp in RvD2 (MRA_1768B, MRA_1768C and mmpL14) were selected and primers were designed. *C*, colony morphology and staining of H37Rv and H37Ra. Cells were harvested and stained with 0.002% neutral red in barbital buffer. *D*, number of viable THP-1 macrophages after infection. *E*, growth of H37Rv and H37Ra in human THP-1 macrophages. * *p* < 0.05.

### Quantitative proteomic characterization of H37Rv and H37Ra

To explore the molecular differences between H37Rv and H37Ra, we compared the proteomic profiles of H37Rv and H37Ra using a label-free quantitative proteomics strategy (**Fig. 2A**). In our assay, a total of 2,724 proteins were identified, covering 67% of the annotated H37Rv proteome (4,032 proteins, TubercuList database) (**Fig. 2B and Supplementary Table 2**). Of these, 2,316 proteins were identified based on the presence of at least two unique peptides and 2,315 proteins were quantifiable. Based on the log_2_-transformed LFQ intensities, we observed very strong correlations among the quantified proteins across replicate samples (R: 0.960 to 0.995) (**Fig. 2C**), suggesting excellent reproducibility of our MS data. Using a stringent cutoff of fold change > 1.5 and *p* < 0.05, we identified 1,209 differentially expressed proteins (DEPs), of which 609 were up-regulated and 600 were down-regulated. A heatmap visualization further highlighted distinct expression patterns between the two strains, reflecting differences in their relative levels of proteins (**Fig. 2D and Supplementary Table 3**).

**Figure 2.**
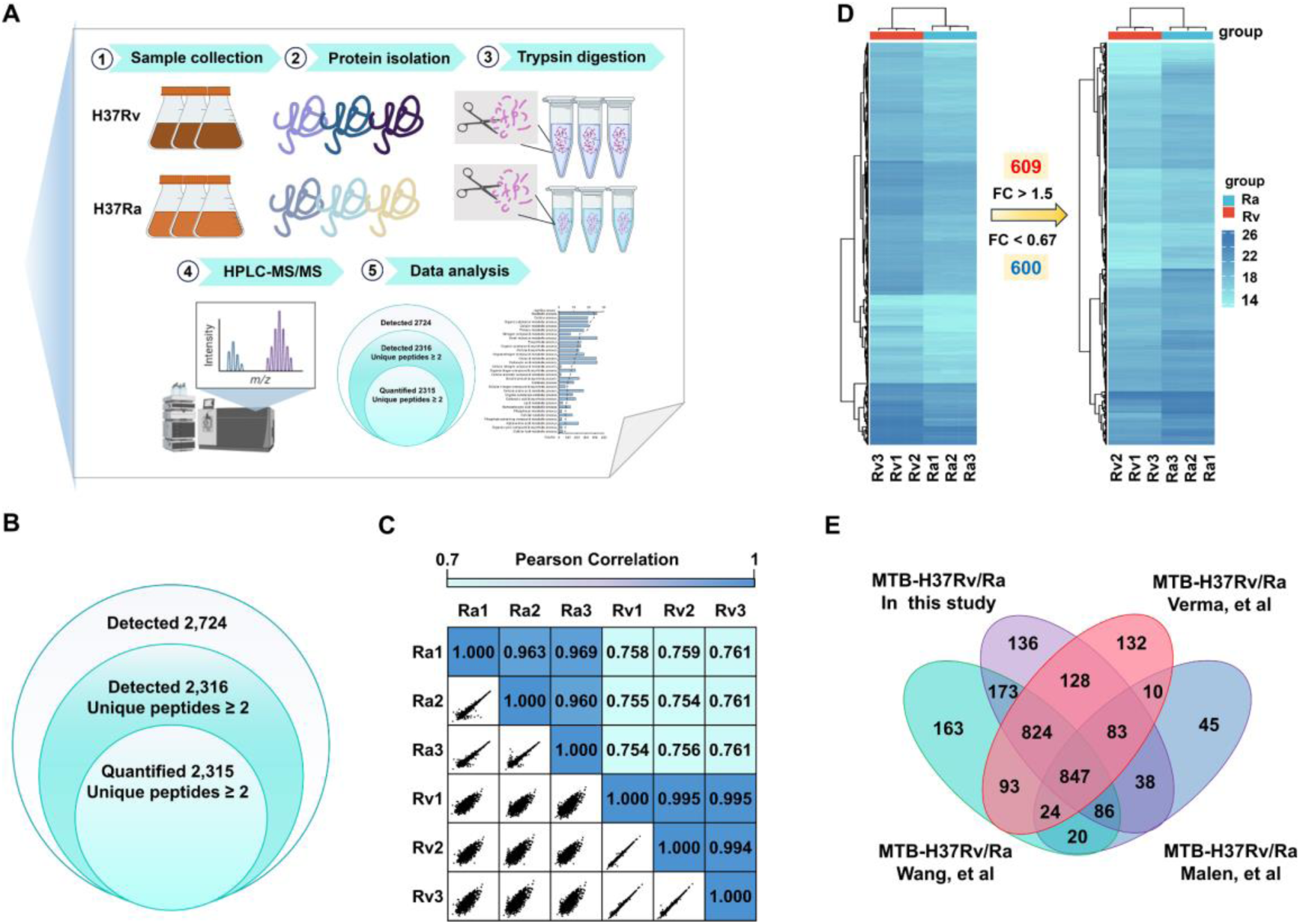
Quantitative proteomics analysis of *M. tuberculosis* H37Rv and H37Ra. *A*, overall workflow of LFQ proteomic strategies. *B*, venn diagram of quantified proteins and differentially expressed proteins. *C*, comparison of LFQ-based quantification intensities within three replicates. *D*, heatmaps of quantified intensity of differentially expressed proteins between H37Rv and H37Ra strains (*p* < 0.05). *E*, comparison of the number of quantified proteins in this study with previous literature.

Using the quantitative protein screen of this study as a criterion, our data were systematically compared and analyzed with the previously published proteomics research results of *M. tuberculosis* ^10–12^. We found that 136 unique proteins in *M. tuberculosis* were quantified in our experiments, and more than 94.1% of the detected proteins were also included in other H37Rv/H37Ra proteomics datasets. Specifically, there were 847 proteins quantified simultaneously in the four datasets, which are likely to be abundant in the strains and play important functional roles (**Fig. 2E and Supplementary Table 4**).

### Functional enrichment analysis of DEPs

To investigate the biological roles of DEPs, we conducted Gene Ontology (GO) enrichment analysis, classifying proteins based on biological processes, molecular functions, and cellular components. In biological process enrichment analysis, up-regulated proteins were enriched in metabolic, cellular, and biosynthesis processes, while down-regulated proteins showed enrichment in bioregulation and macromolecular biosynthesis (Figure 3A-B). For molecular functions, up-regulated proteins focused on catalytic activity, binding (iron/organic heterocycles), and oxidoreductase activity, whereas down-regulated proteins were mainly ribosomal structural components. In cellular localization, up-regulated proteins were in cytoplasm, cell wall, and membrane, while down-regulated proteins localized to membraneless organelle (Figure S3, Supplementary Table 5). KEGG analysis showed up-regulated proteins were involved in purine, β-alanine, and fatty acid metabolism, while down-regulated proteins were enriched in ribosome (Figure 3C-D). These suggested H37Rv prioritized energy metabolism for growth, whereas H37Ra focused on regulatory adaptation and protein synthesis. Notably, catalytic/redox activities are key in lipid metabolism. Lipid enzymes (e.g., fatty acid synthases) localize to cytoplasm, while membrane proteins may mediate phospholipid metabolism/transport. β-alanine is a precursor of pantothenic acid (CoA component), essential for fatty acid metabolism. Thus, H37Rv likely enhances fatty acid metabolism to acquire energy, stabilize biofilms, or evade immunity.

**Figure 3.**
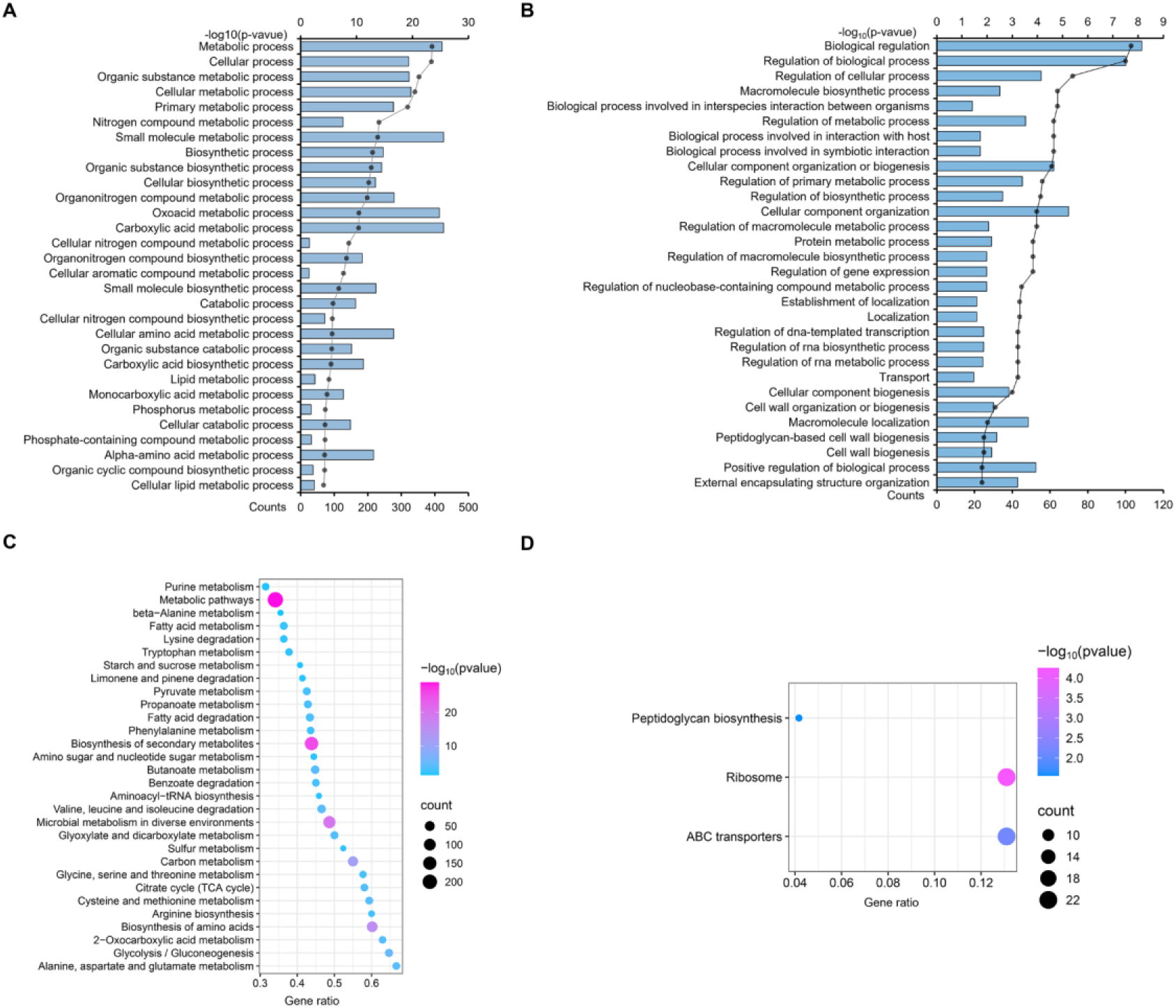
Functional enrichment analysis of DEPs. *A*, GO enrichment analysis of up-regulated DEPs according to biological processes. *B*, GO enrichment analysis of down-regulated DEPs according to biological processes. *C and D*, up- and down-regulated proteins of KEGG enrichment.

### Validation of proteomic data by Western blotting and PRM analysis

To validate our proteomic results, we performed western blotting on seven selected DEPs. The results confirmed the upregulation of known virulence factors, including Rv0251c (Hsp) and Rv3246c (MtrA) in H37Rv. Additionally, proteins involved in fatty acid synthesis, such as Rv3825c (Pks2), Rv1925 (FadD31), Rv3800c (Pks13), and Rv0407 (Fgd1) were also upregulated, while Rv1306 (AtpF) was downregulated. These results were consistent with quantitative proteomics results, confirming the reliability of the proteomics data (**Fig. 4A**). We further validated 177 DEPs through PRM analysis, confirming expression changes for 160 proteins with a fold change > 1.5 and *p* < 0.05 (**Supplementary Table 6)**. As shown in **Figure 4B**, these DEPs were categorized into three functional groups: lipid metabolism, virulence, detoxification, adaptation, and cell wall and cell processes, based on the functional classification of the TubercuList database. Heatmaps demonstrated consistent expression patterns between PRM and quantitative proteomics data. Furthermore, to assess the correlation between PRM-based protein expression and quantitative proteomics data, we performed a Pearson correlation analysis based on the peak area of each peptide in its biological replicates. Based on the scatter plots, the strong linear correlation observed indicated high reproducibility between replicates (**Fig. 4C**). Observe that the expression levels of lipid metabolism-related proteins were significantly elevated in the H37Rv strain, such as KasA, FadD32, and Pks13, which are involved in mycolic acid synthesis ^51^, as well as key proteins in the sulfolipid synthesis pathway, including Pks2 and FadD23 ^52^. Figure 4D shows the extracted ion chromatograms of representative peptides of the Pks2 protein in the H37Rv and H37Ra strains. These findings reveal the important role played by lipid metabolism pathways in the regulation of bacterial virulence.

**Figure 4.**
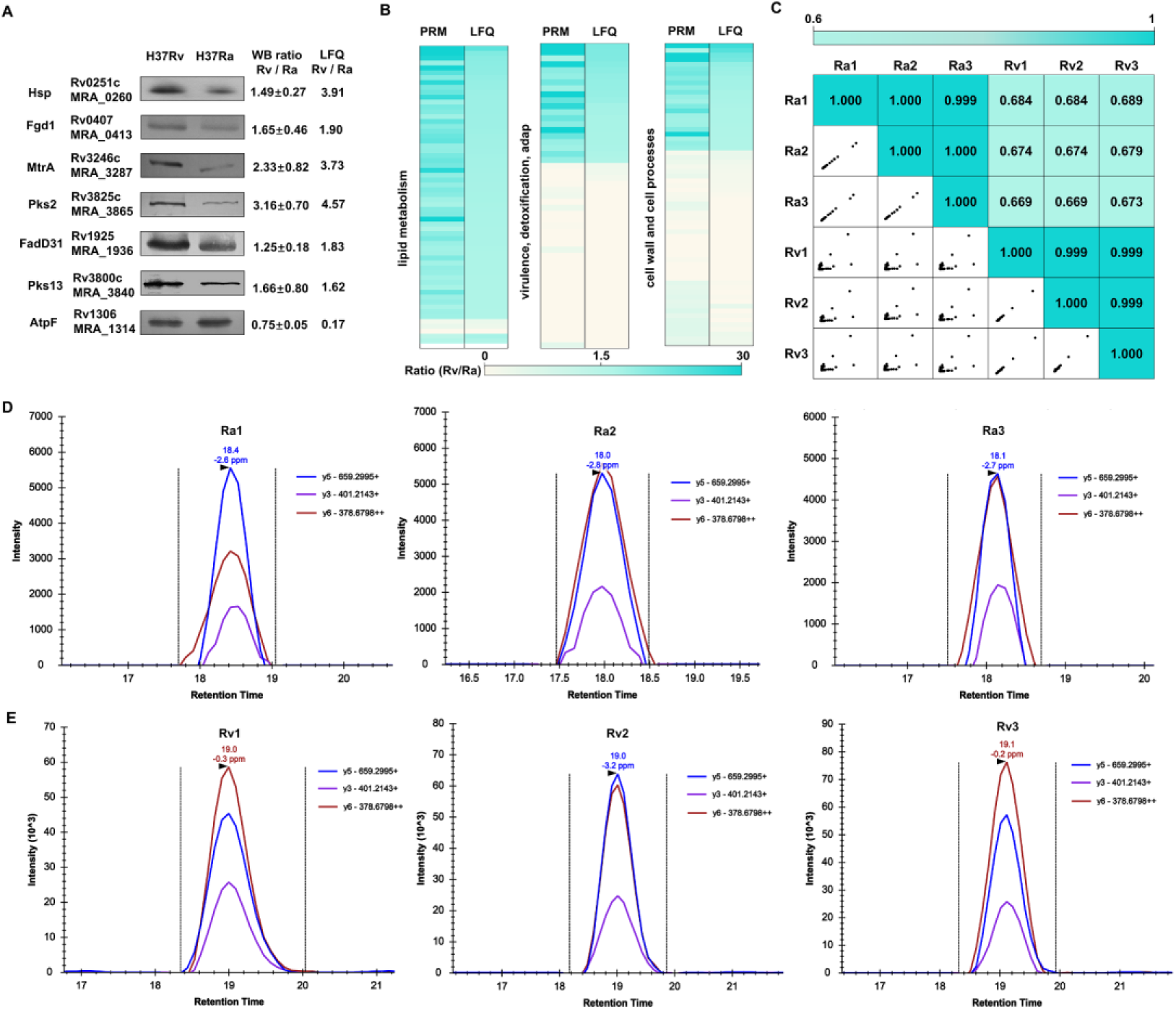
Verification of the proteomic data from H37Rv and H37Ra. *A*, Western blotting analysis of protein expression levels for seven representative DEPs. *B*, heatmaps showing the expression levels of the DEPs selected for validation by PRM. *C*, correlation analysis of converted peak areas for selected DEPs in three groups of biologically replicated samples. *D*, chromatograms of the extracted ion chromatograms (XICs) of representative peptides of the Pks2 protein from strains H37Rv and H37Ra.

### Protein- protein interaction network (PPI)

To further obtain the potential biological roles of the DEPs, a total of 160 PRM-validated DEPs were integrated into the STRING database. We successfully constructed a protein-protein interaction (PPI) network containing 91 proteins with high confidence scores (> 0.7) (**Fig. 5A**). Using the CytoHubba plugin, we identified the most important protein core modules in the network, comprising 20 proteins, namely Rv0860 (FadB), Rv3825c (Pks2), Rv3800c (Pks13), Rv2940c (Mas), Rv1074c (FadA3), Rv0859 (FadA), Rv0131c (FadE1), Rv0154c (FadE2), Rv3140 (FadE23), Rv0400c (FadE7), Rv1070c (EchA8), Rv0675 (EchA5), Rv3039c (EchA17), Rv2245 (KasA), Rv1071c (EchA9), Rv2831 (EchA16), Rv3774 (EchA21), Rv0468 (FadB2), Rv3285 (AccA3), and Rv1323 (FadA4) (**Fig. 5B**). Most of these core proteins are acyl-CoA-related enzymes and polyketide synthases, which are involved in various metabolic pathways, such as fatty acid metabolism and lipid synthesis pathways (**Fig. 5C**). SL-1, an abundant sulfate ester specifically expressed in pathogenic mycobacteria, mainly exists on the outer membrane of *M. tuberculosis*, and its levels positively correlate with the virulence of the strain ^53^. Interestingly, genes involved in the SL-1 biosynthesis pathway *pks2* and *fadD23* showed significant up-regulation in H37Rv, further highlighting their potential role in *M. tuberculosis* virulence.

**Figure 5.**
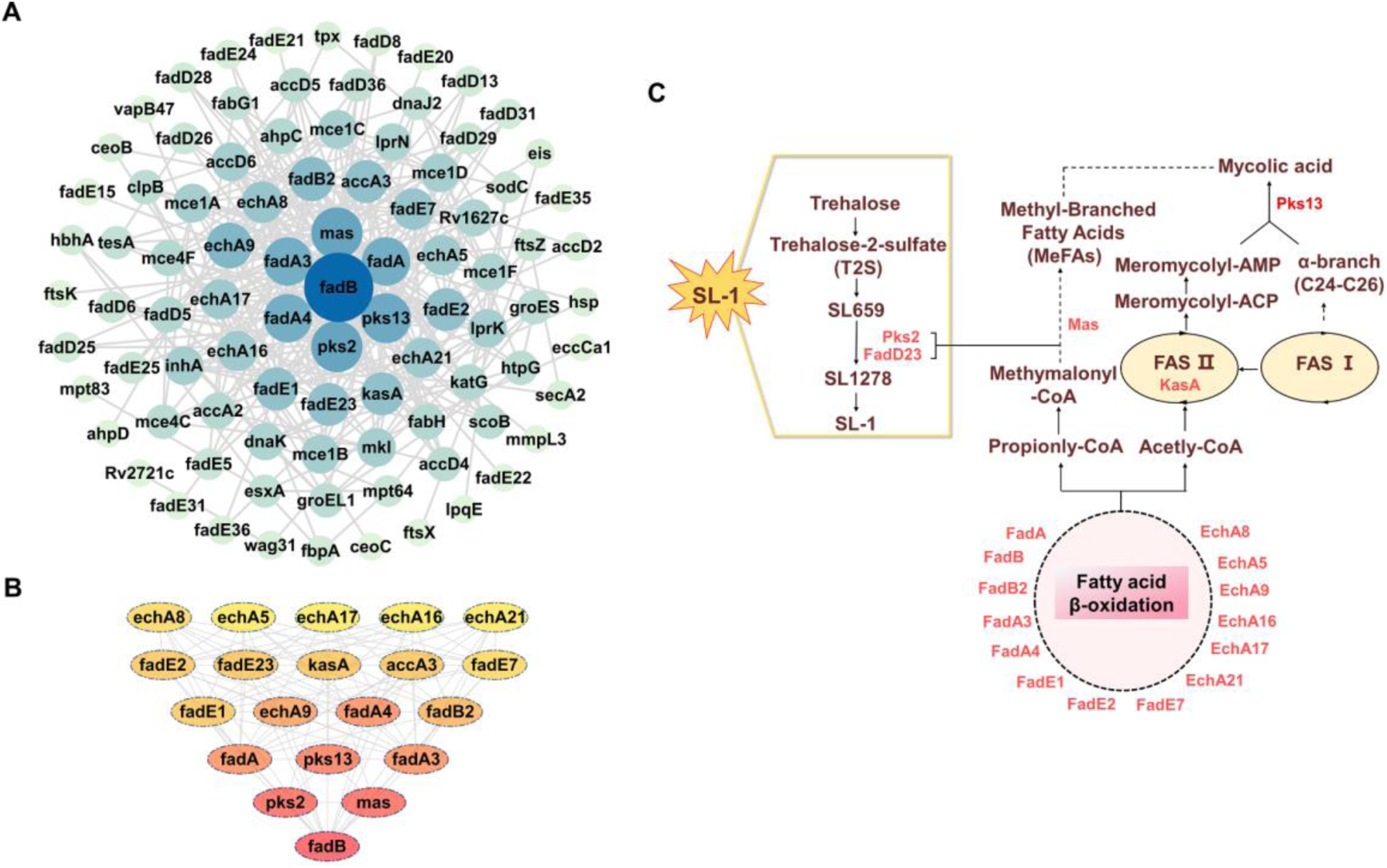
Network analysis of validated DEPs. *A*, visualized PPI network of validated DEPs. *B*, top-ranked network cluster in the PPI network based on Cytohubba plugin (Top 20 DEPs). *C*, illustrations of the 20 differentially expressed proteins involved in lipid and fatty acid metabolism pathways in *M. tuberculosis.* Red shows up-regulated proteins.

### Prediction of virulence associated sRNAs

sRNAs have been found to play key roles in post-transcriptional regulation in a wide range of bacteria, influencing critical cellular processes and helping bacteria in adapting to environmental changes ^54–56^. We hypothesize that ncRNAs targeting potentially DEPs between H37Rv and H37Ra may be involved in the regulation of bacterial virulence. Most sRNAs exert their regulatory effects by base-pairing with target mRNAs, thereby influencing transcription efficiency, mRNA stability, or translation ^27, 57^. In our analysis of non-coding RNAs from H37Rv and H37Ra strains, we identified a total of 3,066 non-coding RNAs in H37Rv. Of these, 2,454 were locatedin antisense strand (AS) regions of genes, while 612 were found in intergenic regions (IGR). For H37Ra, 3,108 non-coding RNAs were predicted, with 2,487 in AS regions and 621 in IGR regions (**Fig. 6A and Supplementary Table 7**). We integrated these predicted sRNAs from H37Rv and H37Ra strains with the validated interaction network of DEPs, suggesting that these proteins could potentially serve as targets for these identified non-coding RNAs, as shown in **Figure 6B**. These proteins were categorized into three main functional groups based on their GO functions listed in the Tuberculist database. Notably, sRNAs appear to be involved in the regulation of various DEPs, with a focus on lipid metabolism, cell wall and cell processes, virulence, adaptation, and detoxification. Interestingly, one gene may be targeted by one or more sRNAs, which exhibit differential expression between H37Rv and H37Ra. For example, the Rv2115c(mpa) was targeted by four different sRNAs, highlighting the complexity of the non-coding RNA regulatory network.

**Figure 6.**
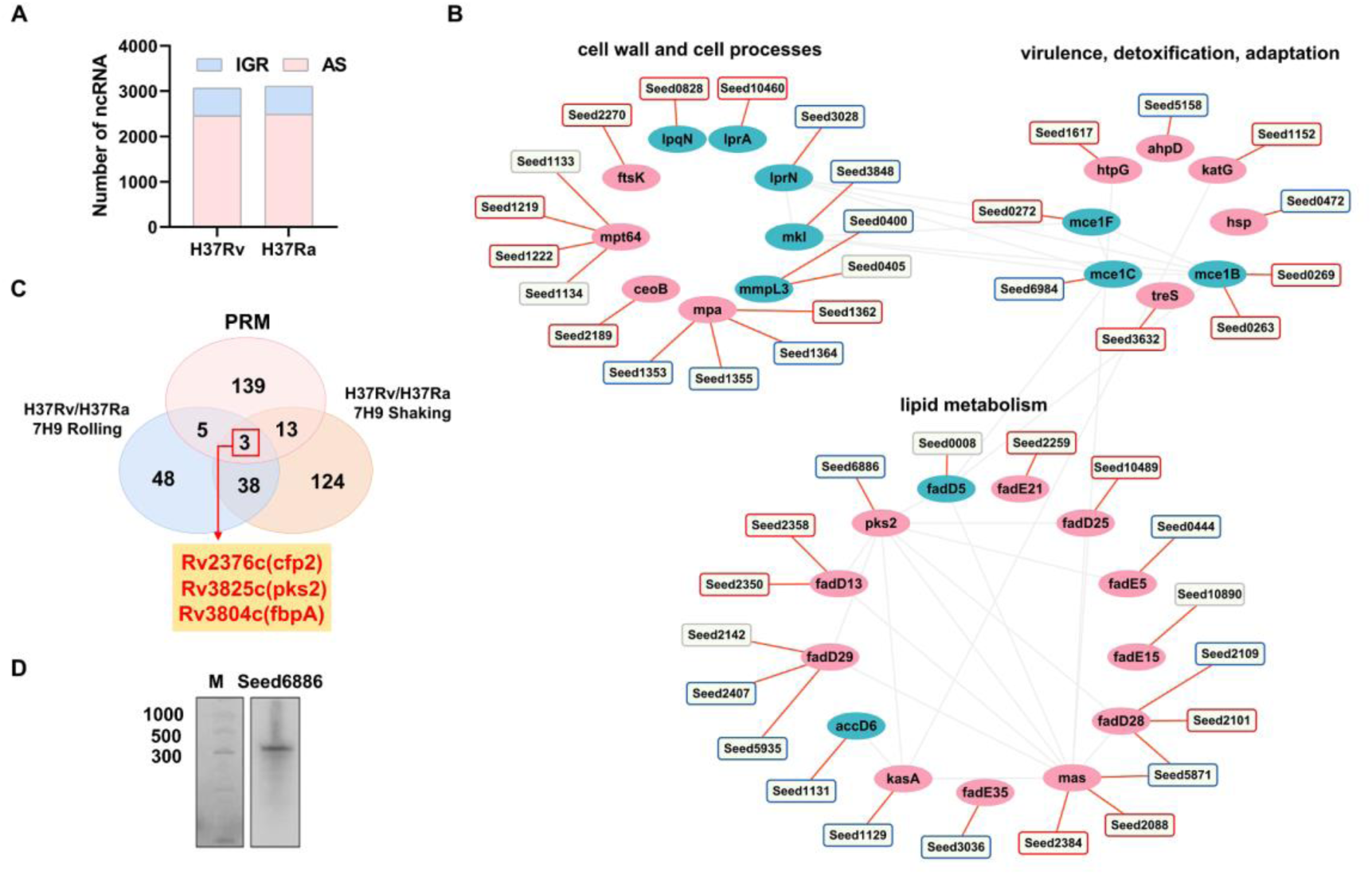
Analysis of sRNAs network. *A*, distribution of sRNAs in *M. tuberculosis* H37Rv and H37Ra. *B*, the network map of sRNAs involved in the regulation of virulence. The squares represent sRNAs and ellipses show proteins, red means “Up”, blue means “Down”, grey means no significance. *C*, overlapping of PRM-validated proteins with differential expressed genes in the GSE3999 dataset. *D*, identification of sRNA presence by Northern blotting.

In an earlier work, Gao *et al.* explored gene expression differences between strains H37Rv and H37Ra under five different nutritional conditions and growth environments ^9^. We analyzed the GSE3999 dataset by using GEO2R, and 41 genes were detected to show a consistent differential expression pattern in 7H9 medium (FC > 1.5 or < 0.67 with adjusted *p*-value < 0.05), regardless of whether using shaking with a ventilated lid or rolling without a ventilated lid (**Supplementary Table 8**). Interestingly, Rv2376c (*cfp2*), Rv3825c (*pks2*) and Rv3804c (*fbpA*) showed significant differential expression in both transcriptome sequencing and PRM analyses (**Fig. 6C**). Using Northern blot, we successfully verified the presence of a novel ncRNA, *Seesd6886*, in strain H37Rv **(Fig. 6D**). Moreover, 3’ RACE analysis was performed by the ligation of an adapter to the 3’ hydroxyl group of H37Rv total RNAs (**Fig. S4**). Given its proximity to *pks2* in the genome, we designated this ncRNA *ASpks2*, the AS RNA of *pks2*.

### Pks2 is the major target of *ASpks2*

As previously described, *pks2* was significantly upregulated at both transcriptional and protein levels in the virulent H37Rv strain. qPCR, Northern blot, and Western blot analyses showed that *ASpks2* transcription was lower in H37Rv compared to H37Ra (Figure 7A & C), whereas pks2 mRNA and protein levels were higher in H37Rv (Figure 7B & D). These results confirmed our hypothesis of a negative regulatory relationship between *ASpks2* and pks2.

**Figure 7.**
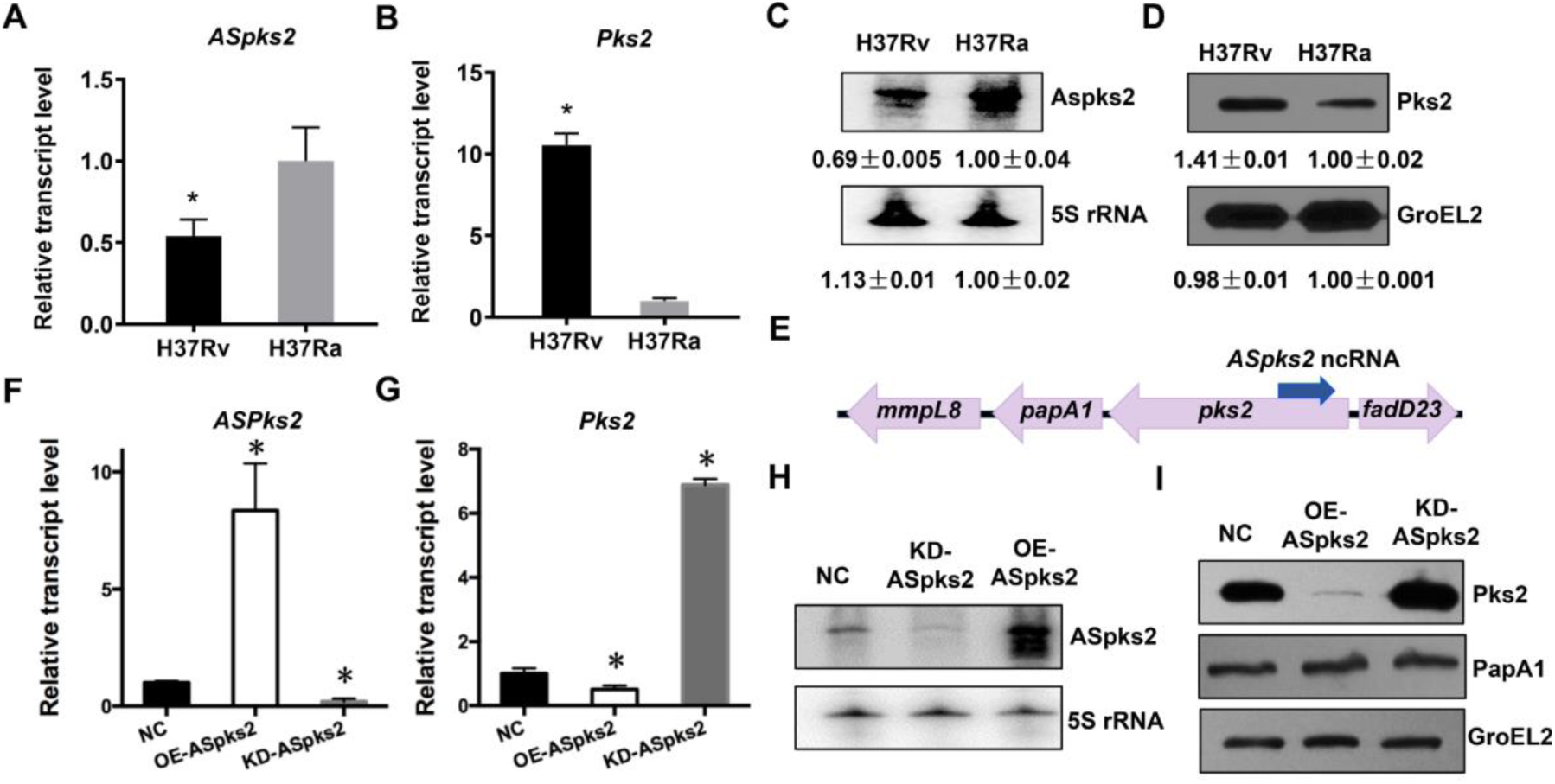
Characterization of *pks2* as the major *ASpks2* target. *A* and *B*, qRT-PCR analysis of *ASpks2* or *pks2* from H37Rv and H37Ra. *C*, northern blot analysis of the *ASpks2* transcript in H37Rv and H37Ra. 5S rRNA was used as a loading control. *D*, western blotting of Pks2 and GroEL2 in the two strains. GroEL2 was used as control for the bacterial integrity of each sample. *E*, schematic representation of downstream or upstream of *pks2* locus. *F* and *G*, qRT-PCR analysis of *ASpks2* and *pks2* in the negative control strain (NC), overexpression strain (OE-*ASpks2*) and knockdown (KD- *ASpks2*) strains. *H*, northern blot analysis of the *ASpks2* transcript in the NC, KD- *ASpks2* and OE-*ASpks2* strains. *I*, western blotting of Pks2, PapA1 and GroEL2 in the three strains (NC, OE-*ASpks2* and KD-*ASpks2*). * *p* < 0.05.

To further determine whether *ASpks2* specifically targets pks2, we constructed the overexpression and knockdown strains of *ASpks2* by cloning the *ASpks2* fragment and its complementary paired fragment into the pMV261 vector. Subsequently, the expression levels of *ASpks2* were verified by qPCR and Northern blotting (**Fig. 7F and H**). In line with our hypothesis, both the mRNA and protein levels of PKS2 were significantly reduced in the OE-*ASpks2* strain, while showing significant upregulation in the KD-*ASpks2* strain (**Fig. 7G and I**). Importantly, the protein levels of PapA1, which is downstream of Pks2, remained unchanged in the OE-*ASpks2* strain or the KD-*ASpks2* strain, confirming the specificity of *ASpks2* in targeting *pks2* (**Fig. 7E and I**). Therefore, *ASpks2* specifically target *pks2*, suppressing its transcriptional level and reducing its protein abundance.

### *ASpks2* promotes H37Rv growth in human macrophages

To explore the biological function of *ASpks2* in *M. tuberculosis* H37Rv, we investigated the phenotypic differences among the control, OE-*ASpks2*, and KD-*ASpks2* strains. In 7H9 liquid medium without rolling, the OE-*ASpks2* strain settled at the bottom of the flask, and the liquid remained clear, while the control and KD-*ASpks2* strains were dispersed evenly throughout the medium. Despite these differences in physical appearance, growth curves for all three strains did not show significant variations (**Fig. 8A-B**). We next investigated the potential role of *ASpks2* in virulence by performing *ex vivo* infections. Human macrophage THP-1 cell lines were infected with negative control, OE-*ASpks2* strain, or KD-*ASpks2* strain, and intracellular growth was assessed. On days 4 and 7 post-infection, the number of viable cells was significantly higher in the OE-*ASpks2* strain compared to the control and KD-*ASpks2* strains (**Fig. 8C**). Colony-forming unit (CFU) counts of bacteria within adherent viable cells revealed a significant increase in colony numbers for the OE-*ASpks2* strain (**Fig. 8D**). *ASpks2* reduced cytotoxicity and necrosis, whereas the negative control and the KD-*ASpks2* strains markedly promoted necrotic cell death, a phenomenon that occurs with the release of the bacterium from lysates during the late stage of infection (**Fig. S5**).

**Figure 8.**
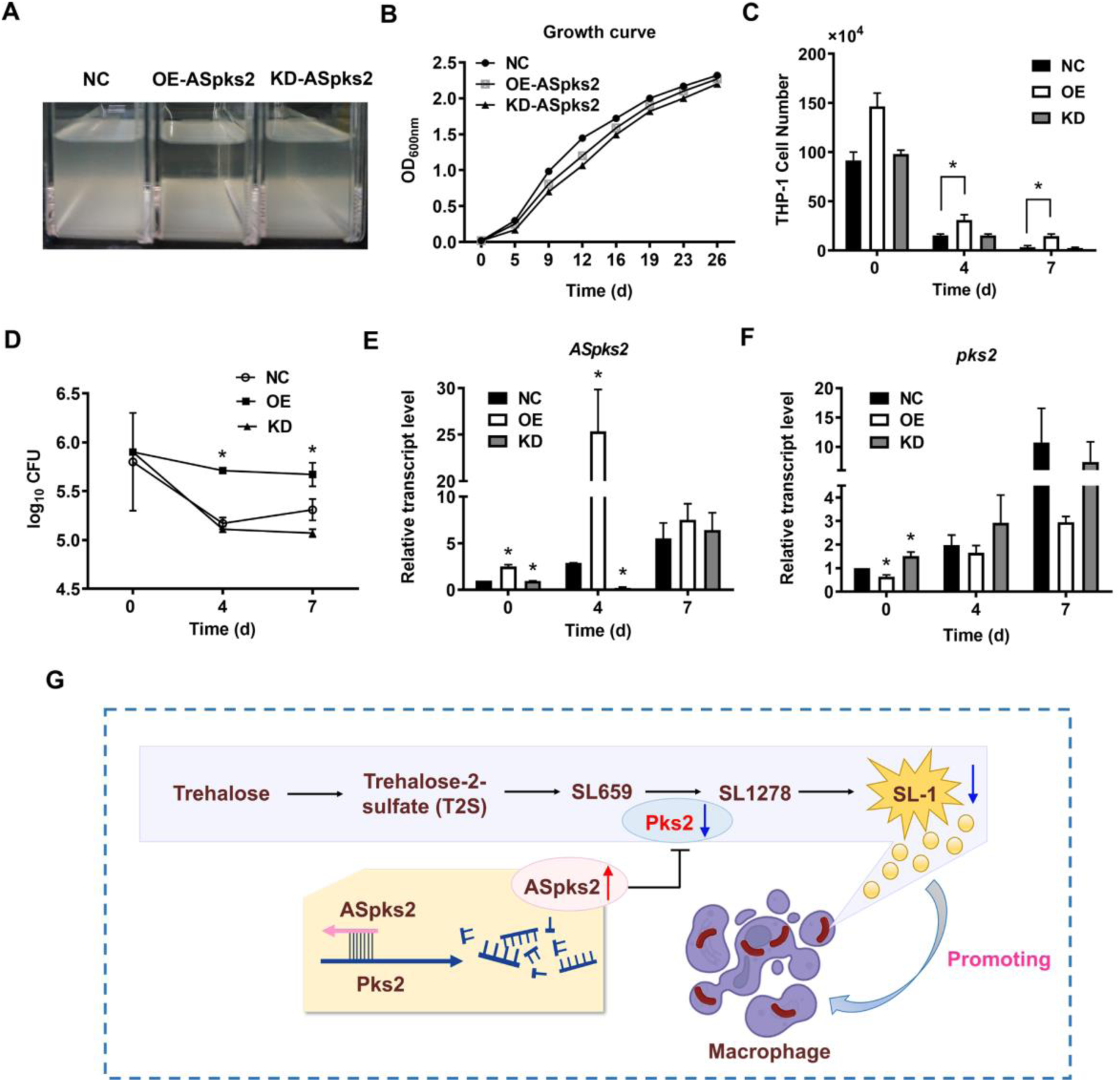
*ASpks2* promotes the growth of H37Rv in human viable THP-1 macrophages. *A*, the three strains were cultured in 7H9 liquid media containing 0.1% Tween 80 till log phase. *B*, growth curves of the NC, OE-*ASpks2* and KD-*ASpks2* strains in 7H9 medium. *C*, viable cell number of THP-1 macrophages after infection. *D*, *ASpks2* promotes intracellular growth of *M. tuberculosis* in human viable macrophages. Human THP-1 macrophages were infected with the indicated strains of *M. tuberculosis.* Adherent macrophages were lysed and plated on solid agar at 4 h (day 0), 4 days and 7 days post-infection, and the number of surviving intracellular bacteria was determined by CFU counting. *E* and *F*, qRT-PCR analysis of *ASpks2* and *pks2* of NC, OE-*ASpks2* and KD-*ASpks2* strains in THP-1 macrophages at 4 h (day 0), 4 days and 7 days post-infection. *G*, potential regulatory role of sRNA *ASpks2* in *M. tuberculosis*. * *p* < 0.05.

Total RNA was extracted from intracellular bacteria after infection, and RT-qPCR was performed to detect *ASpks2* and its target gene, *pks2*. The results showed that *ASpks2* was overexpressed in the OE-*ASpks2* strain during infection, while *pks2* expression was down-regulated (**Fig. 8E-F**). These data indicate that *ASpks2* continues to target *pks2* during infection, thereby enhancing the survival of H37Rv within human macrophages.

Our findings support a model where *ASpks2* mitigates host cell damage by downregulating *pks2* expression. This protective effect may stem from the inhibition of SL-1 synthesis, leading to alterations in the *M. tuberculosis* cell wall composition and contributing to a more stable host-pathogen relationship (**Fig. 8G**). This mechanism highlights the critical role of *ASpks2* in modulating bacterial virulence and enhancing intracellular survival.

## Discussion

sRNAs are critical regulators of bacterial pathogenicity, playing essential roles in processes such as host immune evasion, stress adaptation, and metabolic reprogramming. In this study, we integrated proteomic and transcriptomic analyses to link sRNA regulation to the virulence of *M. tuberculosis*. Within the interaction network of differentially expressed sRNAs and DEPs, we identified a novel sRNA, *ASpks2*, and demonstrate that it negatively regulates the expression of *pks2*, a key enzyme involved in lipid metabolism and virulence in *M. tuberculosis*.

Specifically, *ASpks2* was shown to downregulate *pks2* expression. Overexpression of *ASpks2* led to significant reductions in *pks2* expression, while knockdown of *ASpks2* resulted in its upregulation (**Fig. 8**). Using specific antibodies, we confirmed that *ASpks2* selectively downregulated *pks2* without influencing the protein expression of the downstream gene *papA1*. These findings suggest that *ASpks2* specifically targets *pks2*, influencing the stability of its mRNA and consequently reducing its protein levels. In addition, we found that overexpression of *ASpks2* significantly enhanced the survival of *M. tuberculosis* within human THP-1 macrophages, as evidenced by increased CFU counts and reduced host cell necrosis. *pks2* is involved in the synthesis of SL-1, which is up-regulated during infection of both human macrophages and in mice ^58, 59^. SL-1 has been proposed to play multiple roles in host physiology, including modulation of secretion of pro- and anti-inflammatory cytokines, phagosome maturation arrest, antigen presentation, modulation of membrane dynamics, and host signaling ^60–63^, particularly by suppressing macrophage activation ^64, 65^. While our data do not directly demonstrate that *ASpks2* regulates SL-1 synthesis, we speculate that the downregulation of *pks2* by *ASpks2* may influence SL-1 production. Recent studies have shown that *M. tuberculosis* strains lacking *pks2* exhibit altered SL-1 levels and reduced lysosomal trafficking, which correlates with diminished bacterial clearance and enhanced intracellular survival^65^.

In addition to the role of *ASpks2* in regulating *pks2*, our study underscores the broader impact of sRNAs on *M. tuberculosis* pathogenesis. The integration of proteomic and transcriptomic data revealed a complex network of proteins and pathways involved in the bacterium’s adaptation to the host environment. Specifically, we observed that upregulated proteins in H37Rv were enriched in metabolic process, fatty acid metabolism, cell wall and membrane components. These findings align with the notion that *M. tuberculosis* employs a multifaceted approach to adapt to host immune pressures ^66, 67^, and sRNAs such as *ASpks2* serve as crucial regulators of this adaptive response. By modulating key metabolic and virulence pathways, *ASpks2* enables *M. tuberculosis* to fine-tune its virulence profile and maximize its chances of survival within the host.

Furthermore, this study emphasizes the growing recognition of sRNAs as key players in bacterial adaptation and pathogenesis. Similar regulatory roles for sRNAs have been identified in other pathogens, such as *Pseudomonas aeruginosa*, *Staphylococcus aureus* and *Listeria*, where sRNAs influence both chronic and acute infection regulation, as well as immune evasion ^68–70^. The conservation of sRNA-mediated regulatory mechanisms across different pathogens suggests that targeting sRNA networks could offer a promising therapeutic strategy for modulating bacterial virulence. Disrupting sRNA regulation could reduce the pathogen’s ability to adapt to the host and enhance immune responses, providing a potential complement to traditional antibiotics, particularly in the context of drug-resistant strains. By targeting multiple virulence factors, this strategy could make it easier for the host immune system to clear the infection.

Although this study provides valuable insights into the role of *ASpks2* in *M. tuberculosis* virulence, several key areas require further investigation. First, while we have shown that *ASpks2* downregulates *pks2*, future work should focus on validating the impact of *ASpks2* on SL-1 synthesis through lipidomic analysis. Second, further studies are needed to explore how *ASpks2* interacts with other sRNAs and regulatory proteins to coordinate the broader regulatory network that governs *M. tuberculosis* virulence. Understanding the full scope of regulatory actions of *ASpks2* will provide a more comprehensive understanding of its role in bacterial pathogenesis. Lastly, investigating the role of *ASpks2* under stress conditions, such as nutrient limitation, hypoxia, and antibiotic exposure, will shed light on its contribution to *M. tuberculosis* survival in latent and chronic infections.

In conclusion, the integrated transcriptomic and proteomic analysis establishes a sRNA regulatory network in *M. tuberculosis*. We identify a new sRNA *ASpks2* that targets *pks2* and affects the survival of *M. tuberculosis* in human macrophage cells. This study serves as the first proteome-wide analysis of sRNA regulatory networks and provides new clues for the mechanism involved in virulence of *M. tuberculosis*.

## Supporting information

All Supplemental Figures

## ACKNOWLEDGEMENT

This work was supported by the National Key Research and Development Program of China (2023YFC2307202), the National Natural Science Foundation of China (32394014 & 82304575), and Major Project of Guangzhou National Laboratory (GZNL2024A01023). The authors thank Min Wang (Analysis and Testing Center, Institute of Hydrobiology, CAS) for the assistance with the proteomic experiments.

## DATA AVAILABILITY

RNA-seq data were deposited in the National Center for Biotechnology Information, under accession number PRJNA1196512 (https://www.ncbi.nlm.nih.gov/bioproject/?term=PRJNA1196512). The raw MS data have been deposited to the public access ipro x database (http://www.iprox.org) with the identifier PXD064766.

## AUTHOR CONTRIBUTIONS

F.G., L.B., and Y.C. conceived and designed the experiments and wrote the article. Q.W., Y.C., and M.Y performed the experiments. W.Y., L.Y., F.J., Y.C. and L.B. analyzed the data. All authors read and approved the final article.

## SUPPLEMENTAL DATA

This article contains supplemental data.

## Abbreviations

M. tuberculosis: Mycobacterium tuberculosis
sRNA: small RNA
*ASpks2*: antisense stranded small RNA of *pks2*
DEPs: differentially expressed proteins
LFQ: label free quantification
LC-MS/MS: liquid chromatography-tandem mass spectrometry
MS/MS: tandem mass spectrometry
DIA: data-independent acquisition
AUC: area under the curve
PRM: parallel reaction monitoring
PBS: phosphate buffer saline
FA: formic acid
PVDF: polyvinylidene fluoride
FC: fold change
FDR: false discovery rate
GO: Gene ontology
KEGG: Kyoto Encyclopedia of Genes and Genomes
AS: antisense stranded
IGR: intergenic region
CFU: colony-forming unit
MOI: multiplicity of infection
SD: standard deviation
SL-1: Sulfolipid-1.

