## Supplementary material for "Small RNA *ASpks2* Promote *Mycobacterium tuberculosis* Survival in Macrophages *via* Targeting Polyketide Synthase 2": All Supplemental Figures

**Supplementary Figures**

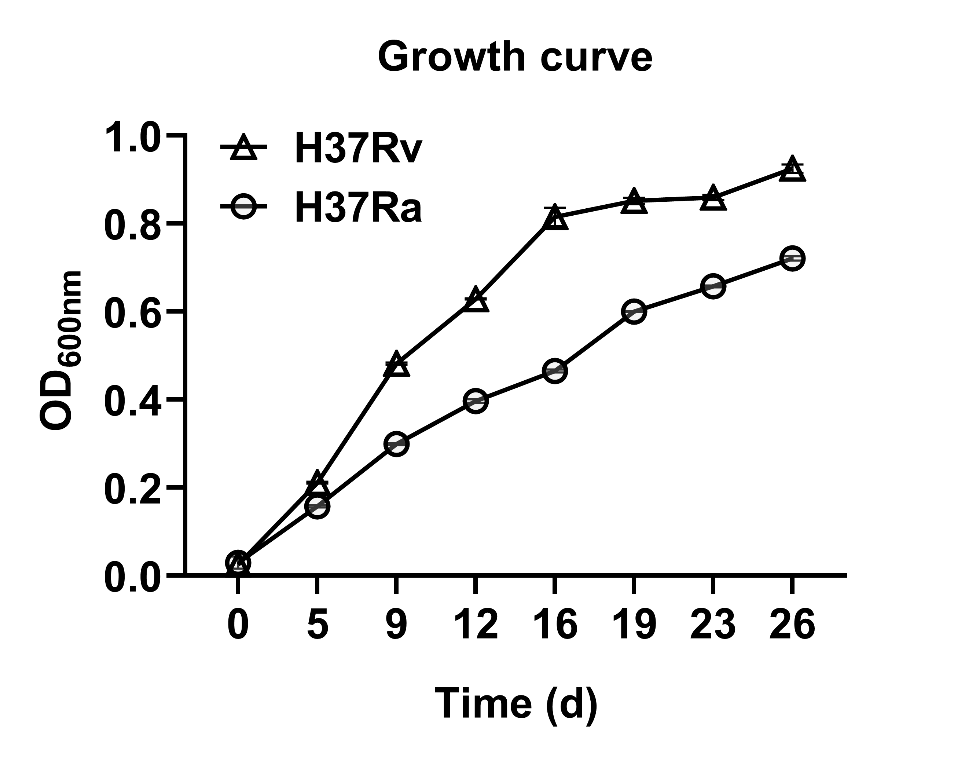

**Figure S1.** Growth curves of *M. tuberculosis* H37Rv and H37Ra in 7H9 medium (10% ADC + 0.5% glycerol + 0.05% Tween 80).

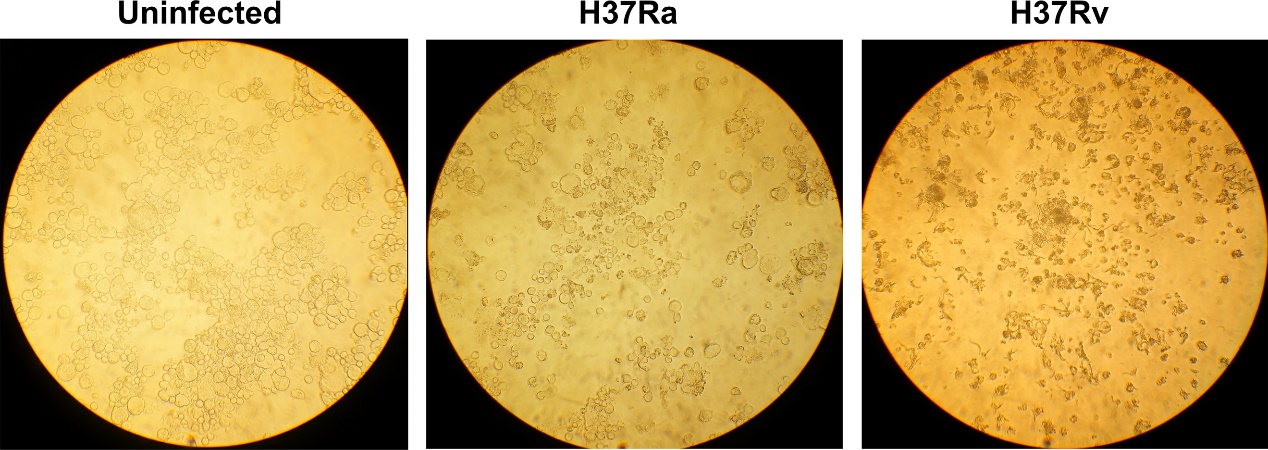

**Figure S2**. Morphologic changes in THP-1 macrophages after infection by *M. tuberculosis*.

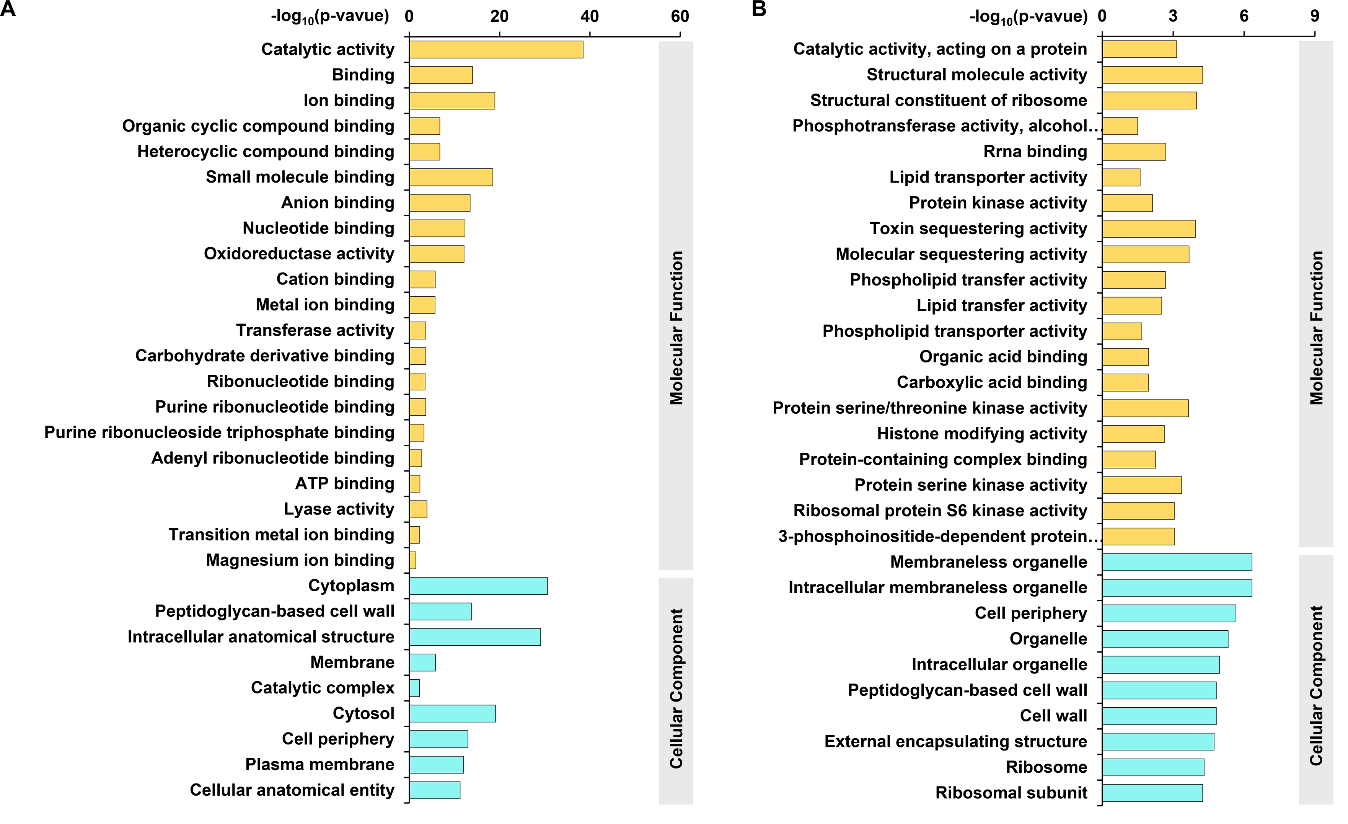
**Figure S3.** GO enrichment analysis of differentially expressed proteins based on cellular components and molecular functions. *A,* up-regulated *B*, down-regulated.

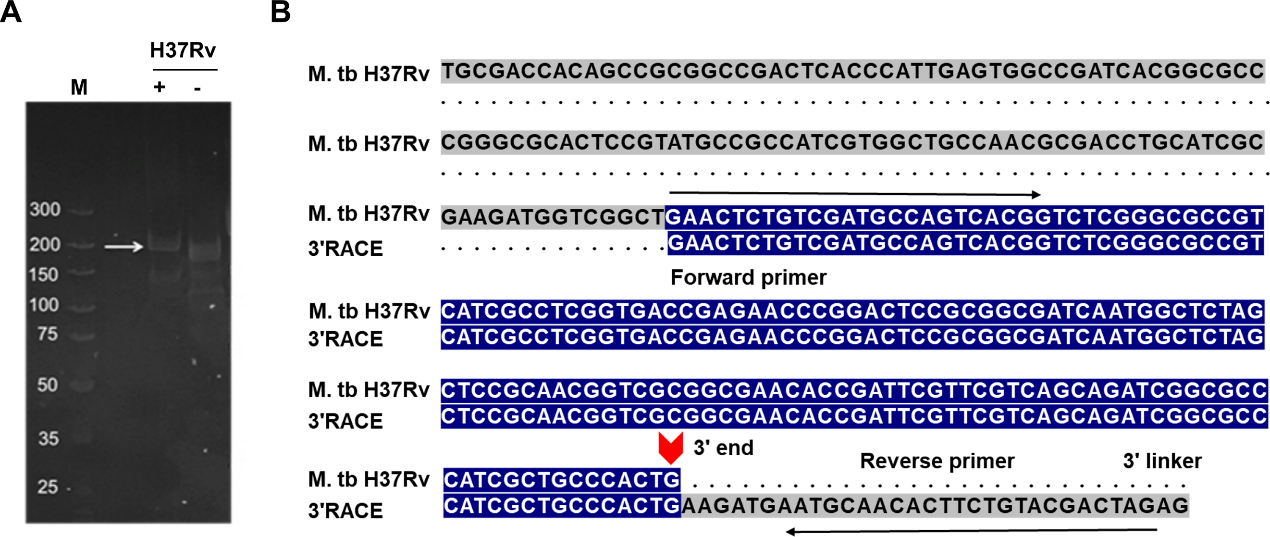

**Figure S4.** Verification of *ASpks2* by 3' RACE. *A*, 3ʹ RACE amplification of *ASpks2* 3' end. M: DNA molecular weight marker (bp); +, reverse transcription in the presence of reverse transcriptase; -, reverse transcription in the absence of reverse transcriptase. *B,* schematic representation of the *ASpks2* 3ʹ end.

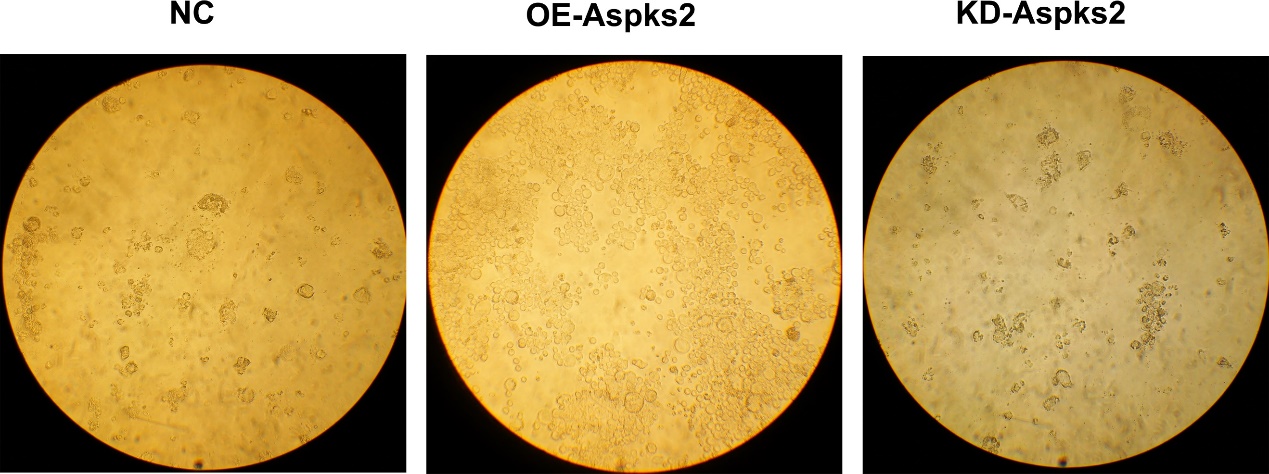

**Figure S5.** Morphological changes of THP-1 macrophages under different conditions of infection.

**Supplementary Table 1.** List of primers used in this study.

**Supplementary Table 2.** List of proteins quantified by LC-MS/MS analysis of *M. tuberculosis* strains H37Rv and H37Ra at log-phase.

**Supplementary Table 3.** List of all quantified differentially expressed proteins in *M. tuberculosis* H37Rv and H37Ra.

**Supplementary Table 4.** Comparison of the quantified proteins of H37Rv and H37Ra in log-phase with Wang et al, Verma et al and Malen et al.

**Supplementary Table 5.** Complete list of GO terms and KEGG pathways enriched by differential proteins in *M. tuberculosis* H37Rv and H37Ra.

**Supplementary Table 6.** Target proteins quantification in PRM experiment.

**Supplementary Table 7.** Small RNA sequencing analysis of H37Rv and H37Ra in *M. tuberculosis*.

**Supplementary Table 8.** Differential expression analysis of the GSE3999 dataset.
